# A bacterial resolvase first exploits, then constrains intrinsic dynamics of the Holliday junction to direct recombination

**DOI:** 10.1101/644575

**Authors:** Sujay Ray, Nibedita Pal, Nils G. Walter

**Affiliations:** Single Molecule Analysis Group, Department of Chemistry, University of Michigan Ann Arbor, Michigan 48109, USA

**Author notes:** These authors contributed equally (Sujay Ray, Nibedita Pal). Indian Institute of Science Education and Research (IISER) Tirupati, Tirupati 517507, Andhra Pradesh, India (Nibedita Pal).

## Abstract

Homologous recombination forms and resolves an entangled DNA Holliday Junction (HJ) critical for achieving genome repair. We use single-molecule observation and cluster analysis to probe how prototypic bacterial resolvase RuvC selects two of four possible HJ strands for cleavage. RuvC first exploits, then constrains intrinsic HJ isomer exchange and branch migration dynamics to direct cleavage toward only a desired, catalytically competent HJ conformation, thus controlling recombination products.

---

In Gram-negative bacteria such as *Escherichia coli* (*E. coli*), Holliday Junction (HJ) resolution is achieved by endonuclease RuvC, a well-studied, prototypical resolvase with extremely high topology and sequence specificity^1–3^. To achieve site-specific resolution, the HJ needs to be correctly positioned through branch migration of the junction, which occurs either spontaneously or catalyzed by the RuvAB complexd^4,5^. A homodimer of RuvC can bind a HJ independently of RuvAB^6–8^ to distort the junction at the crossover point, eventually introducing two interdependent symmetric nicks via a two-Mg^2+^ catalytic mechanism^2,9,10^. Numerous *in vitro* studies with synthetic DNA have shown that RuvC recognizes a HJ structurally, but in sequence-independent manner. More specifically, in a recent study Zhou *et al.*^11^ showed that a RuvC-bound HJ retains its intrinsic fluctuations^12–15^ between two alternatively stacked, X-shaped conformational isomers (*iso-I* and *iso-II*), while an intermittent, multivalently bound, yet partially dissociated (PD) RuvC allows the HJ to undergo nearly unimpeded conformer exchange as well as branch migration^11^. However, it has remained unclear how RuvC achieves its remarkable cleavage specificity for the cognate 5’-ATT^↓^X-3’ sequence (X = G/C; ^↓^= cleavage site) to ensure proper control over chromosome segregation^1,3,16–18^.

Here we leverage single-molecule Förster resonance energy transfer (smFRET) to investigate the dynamic interaction of *E. coli* RuvC with a HJ that leads to sequence-specific junction resolution. Our results reveal that the resolvase binds to both cleavage-incompetent (*iso-I*) and –competent (*iso-II*) conformations, wherein the cognate sequences reside on the two bent and continuous (i.e., non-crossover) strands, respectively (Fig. 1a). Critically, RuvC takes advantage of the intrinsic HJ conformational and branch migration dynamics to find and then kinetically trap the catalytically active conformation through a (near-)irreversible switch from fluctuations between *iso-I* and a partially open state to fluctuations between *iso-II* and the partially open state. On the basis of these findings, we propose a model wherein RuvC exploits the intrinsic HJ dynamics until the cognate sequence enters the active site, inducing a snap-lock conformational switch that helps RuvC achieve high sequence specificity at little energetic cost.

**Figure 1.**
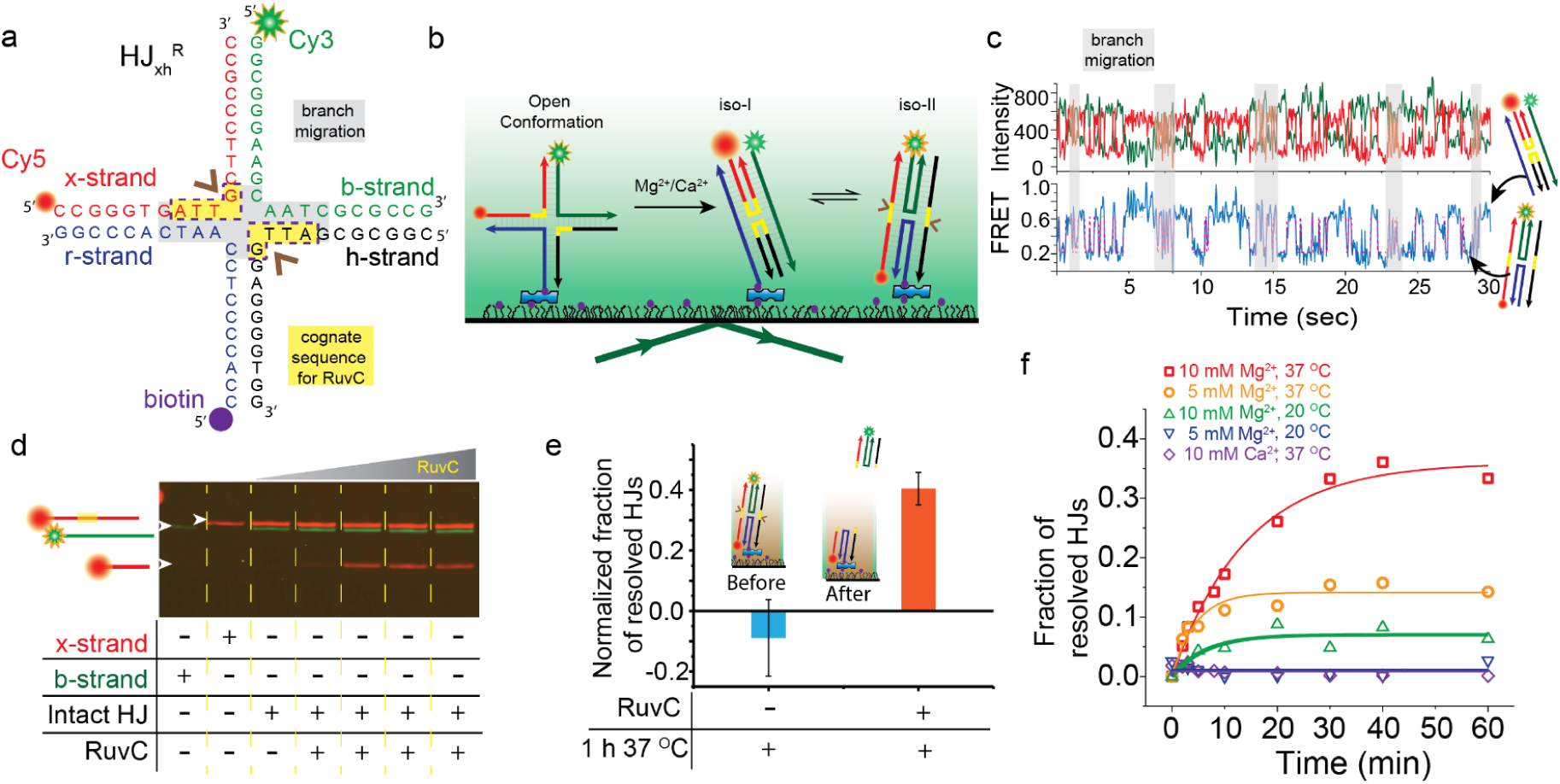
Holliday junction assay design and cleavage by RuvC. **a,** Schematic and sequence of the HJ_xh_^R^ design. The nucleotides highlighted in yellow represent the cognate cleavage sequence for RuvC. The nucleotides highlighted in gray represent the region of the junction that can potentially branch migrate due to sequence homology. The brown arrows indicate the cleavage sites. The 5’ ends of the r, x, and b-strand are labeled with biotin, Cy5 and Cy3, respectively. **b,** Schematic of our smFRET assay under TIRF illumination to monitor the dynamics of HJ_xh_^R^ molecules. The HJ transitions between two isomeric states *iso-I* and *iso-II*. The cognate sequences are shown in yellow in the bent, cleavage-incompetent *iso-I* and continuous, cleavage-competent *iso-II* strands. **c,** Representative single-molecule time trajectory showing donor (green) and acceptor (red) intensity in the top panel, FRET (blue) and HMM fitting (magenta) in the bottom panel. Fast transitions highlighted in gray are attributed to branch migration dynamics. Representative cartoons indicate the conformational states associated with particular FRET values. **d,** Superimposed Cy3 (green) and Cy5 (red) scans of a denaturing urea-polyacrylamide gel showing RuvC-mediated cleavage of HJ_xh_^R^ in the presence of varying concentrations of RuvC. The top bands represent the full-length 20-nt-long DNA; the bottom bands are the cleaved 10-nt-long DNA product. The schematics on the right-hand side represent the relative length of DNA corresponding to each band. The Cy3-labeled b strand is not cleaved due to the absence of a cleavage sequence. **e,** Quantification of cleavage using a single molecule assay. Normalized ratios of number of colocalized Cy3-Cy5 spots and number of all Cy5 spots before and after cleavage are plotted for the presence and absence of RuvC. Without RuvC, this ratio remains close to zero. In the presence of 10 mM Mg^2+^ and 400 nM RuvC incubated at 37°C for 1 h, this ratio becomes 0.40±0.05, representing RuvC-mediated cleavage. The inset shows a schematic of the experiment before and after cleavage in the presence of RuvC. **f,** Quantitative analysis of a time course of RuvC-mediated cleavage of HJ_xh_^R^ in the presence of 10 mM Mg^2+^ at 37 °C (red square), 5 mM Mg^2+^ at 37 °C (orange circle), 10 mM Mg^2+^ at 20 °C (green up triangle), 5 mM Mg^2+^ at 20 °C (blue down triangle), and 10 mM Ca^2+^ at 37 °C (purple diamond) at 400 nM RuvC. Saturation curves are fit to the data. The fraction of HJ_xh_^R^ molecules cleaved by RuvC decreases with decreasing temperature and decreasing Mg^2+^ concentration. This provides us with an opportunity to decouple conformational dynamics from cleavage.

To explore RuvC mediated cleavage, we chose an HJ sequence that contains the resolvase’s cognate sequence within 5 base pairs of sequence homology and positioned for optimal cleavage activity^16^ across the junction on the x- and h-strands (HJ_xh_ ^R^, Fig. 1a). Cyanine-3 (Cy3, donor) and Cyanine-5 (Cy5, acceptor) dyes on two arms of the HJ allow monitoring the conformational dynamics using total internal reflection fluorescence (TIRF) based smFRET (Fig. 1b), as exploited for other HJs (Supplementary Fig. 1)^14,15,19–21^. As expected, in the presence of divalent metal ions Mg^2+^ or Ca^2+^ HJ_xh_ ^R^ undergoes fast dynamic transitions between stacked isomeric conformers^14,15,19–21^ *iso-I* and *iso-II* of high (E_FRET_ = 0.65±0.12) and low (E_FRET_ = 0.21±0.11) FRET efficiencies, respectively (Fig. 1c). The interconversion rate constants of k_*I→II*_ = 2.8±0.3 s^-1^ and k_*II→I*_ = 4.0±0.4 s^-1^ at 5 mM Mg^2+^ decrease gradually with increasing Mg^2+^ concentration (Supplementary Table 1; Supplementary Fig. 2). Occasional branch migration of the HJ was detected as a variation in FRET dynamics by visual inspection, as observed before^11^. A Fano-factor of >1, evaluating non-randomness through the variability in number of pairwise transitions relative to its mean^22,23^, was found to be a signature of the co-existence of regimes of slow and fast FRET transitions representing *iso-I*↔*iso-II* isomerization and branch migration, respectively (Fig. 1c and Supplementary Fig. 3). When Mg^2+^ was replaced with Ca^2+^, only slightly slower kinetics were observed, indicating that HJ_xh_^R^ shows similar two-state isomerization dynamics in Mg^2+^ or Ca^2+^ as other HJs (Supplementary Fig. 1)^14,15,19–21^, with a slight (60%:40%) *iso-I*:*iso-II* bias dictated largely by the junction sequence^19,24^.

The HJ_xh_ ^R^ cleavage activity of RuvC was tested upon incubation for 1 h at 37 °C in standard cleavage buffer (20 mM Tris-HCl, pH-8, 20 mM NaCl, 1 mM EDTA, 1mM DTT, 0.1mg/ml BSA, 10 mM MgCl_2_) by monitoring the appearance of the shortened Cy5 labeled cleavage product using denaturing gel electrophoresis (Fig. 1d). Under these conditions, increasing the RuvC concentration yielded an isotherm with a half-saturation value *K*_1/2_ of ∼66 nM RuvC (Supplementary Fig. 4). In our TIRF microscope, we instead monitored the disappearance of the no longer surface-coupled Cy3-labeled product spots (Fig. 1e), confirming that RuvC resolves single HJ_xh_ ^R^ molecules at 37 °C in standard cleavage buffer. At a saturating concentration of RuvC (400 nM), the cleavage rate constant was 1.58 × 10^-3^ s^-1^, similar to previous studies^16^, and became zero when Mg^2+^ was replaced with Ca^2+^ (Fig. 1f). Bulk cleavage reactions performed at room temperature and/or only 5 mM Mg^2+^ showed significantly reduced cleavage (Fig. 1f), as expected^25^, providing an opportunity to decouple the impact of RuvC on HJ_xh_ ^R^ conformational dynamics observed by smFRET from HJ cleavage (which is ∼35-fold less efficient at our standard decoupled smFRET conditions of 20 °C in 5 mM Mg^2+^ compared to 37 °C in 10 mM Mg^2+^, Fig. 1f).

Using decoupled smFRET conditions in the presence of saturating (400 nM) RuvC, we observed that the dynamic exchange between *iso-I* and *iso-II* becomes more complex (compare Figs. 2a and 2b); in particular, a new mid-FRET state appears (E_FRET_ = 0.4±0.1) that lies between the high-FRET *iso-I* (E_FRET_ = 0.64±0.13) and the low-FRET *iso-II* (E_FRET_ = 0.17±0.1). This state is similar to the planar open conformation of the HJ in the absence of divalent metal ions^10,11^, and is henceforth termed “open”, or Op. Upon replacing Mg^2+^ with Ca^2+^, this Op state disappears, leaving only *iso-I* (E_FRET_ = 0.68±0.09) and *iso-II* (E_FRET_ = 0.15±0.08, Fig. 2c,d). The observation of Op only in the presence of RuvC and Mg^2+^, which are both required for cleavage, suggests that this state is relevant for HJ resolution. In fact, visual inspection revealed that ∼40% of smFRET trajectories visit Op, while the remainder continues to show two-state behavior only, most likely due to limited RuvC binding^16^ (Fig. 2e). Notably, a similar fraction (∼37%) of HJ molecules is cleaved under optimal cleavage conditions (Fig. 1f), further supporting the notion that the population accessing the Op state under decoupled smFRET conditions is RuvC-bound and cleavage-enabled.

**Figure 2.**
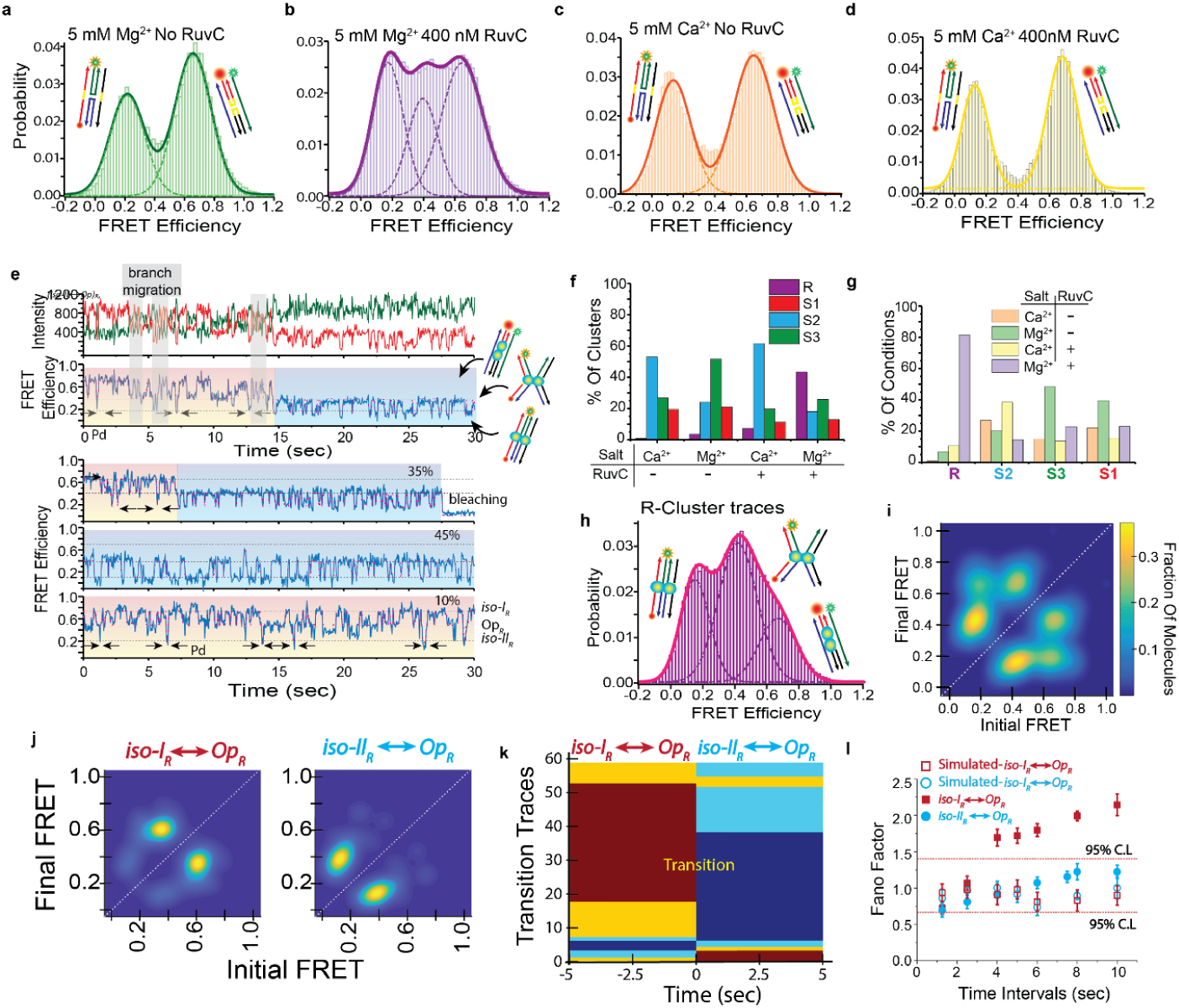
Single molecule Cluster Analysis (SiMCAn) of smFRET trajectories reveals dynamic behaviors unique to the cleavage-competent HJ-RuvC complex. FRET probability distributions of **a,** 5 mM Mg^2+^ without RuvC, **b,** 5 mM Mg^2+^ plus 400nM RuvC **c,** 5 mM Ca^2+^ without RuvC, and **d,** 5 mM Ca^2+^ plus 400nM RuvC. Multi-peak Gaussian functions are used to fit each histogram. Associated structural cartoons depict the conformational states corresponding to each FRET population. **e,** Representative fluorescence intensity trace of Cy3 (green) and Cy5 (red) fluorophore, FRET efficiency (blue) and HMM fitting (magenta) from the 5 mM Mg^2+^plus 400nM RuvC condition (top panel). Three additional FRET efficiency traces show different types of behaviors under the same conditions (bottom panel), with percentages delineating the fractions of traces displaying such behavior. Associated cartoons represent most likely conformations corresponding to each FRET value. The orange background represents *iso-I*_*R*_ *↔ Op*_*R*_ behavior and the blue background represents *iso-II*_*R*_ *↔ Op*_*R*_ behavior. Sudden visits to non-prevalent FRET states, assigned as RuvC’s PD state (marked by black arrows) and branch migration (marked by gray background), are observed in the *iso-I*_*R*_ *↔ Op*_*R*_ behavior and absent from the *iso-II*_*R*_ *↔ Op*_*R*_ behavior. **f,** Summary of SiMCAn results, showing bar graphs with occupancy of the four clusters found in four experimental conditions. Clusters S1, S2 and S3 are distributed over all different conditions. By contrast, the R-cluster is predominantly found in the 5mM Mg^2+^ plus 400 nM RuvC condition, indicating that the R-cluster is associated with the RuvC-HJ_xh_^R^ complex. **g,** Bar graph showing the fraction of molecules from of each experimental condition contributing to the four clusters. More than 80% of the molecules in the R-cluster belong to the condition where both Mg^2+^ and RuvC are present. **h,** FRET efficiency histogram calculated from the traces belonging to the R-cluster, shown with multi-peak Gaussian fits. Associated cartoons represent different RuvC-HJ complex conformations corresponding to different FRET states. **i,** Transition Occupancy Density Plots (TODPs) from R-cluster traces showing the fraction of HJs that undergo conformational transitions from a given initial FRET state to a specific final FRET. Three main sets of bidirectional transitions are observed, representing *iso-II*_*R*_ *↔ Op*_*R*_, *iso-I*_*R*_ *↔ iso-II*_*R*_, and *iso-I*_*R*_ *↔ Op*_*R*_ behavior, respectively. **j,** TODP calculated from the early phases of the traces (red/yellow) show only *(iso-I ↔ Op)*_*R*_ behavior (left panel), whereas segments from the later phases of the traces (cyan/blue) show *iso-II*_*R*_ *↔ Op*_*R*_ behavior (right panel). **k,** The early and late phases of the mixed R-cluster traces were classified into four clusters (annotated in red, yellow, cyan and blue) using SiMCAn. The early phases, representing *iso-I*_*R*_ *↔ Op*_*R*_ behavior, and the late phases, representing *iso-II*_*R*_ *↔ Op*_*R*_ behavior, belongs to different clusters (see Supplementary Fig.7a). Upon reconstitution of their early and late phases with their clusters annotated, a clear propensity becomes evident for individual traces to converge over time onto the *iso-II*_*R*_ *↔ Op*_*R*_ behavior. **l,** The Fano factor was calculated across various time intervals for the *iso-I*_*R*_ *↔ Op*_*R*_ behavior (solid red square), *iso-II*_*R*_ *↔ Op*_*R*_ behavior (solid blue circle) and simulated Poisson data same in length and number of traces for the same behaviors (open red square, open blue circle) respectively. The dashed lines indicate the 95% confidence level of the data. The *iso-II*_*R*_ *↔ Op*_*R*_ behavior Fano factor values deviate from 1, indicating a non-random underlying distribution, while the *iso-II*-loaded behavior Fano factor data remain close to 1, indicating a Poisson distribution.

To classify the dynamic features of these RuvC-HJ complexes, we utilized model-independent hierarchical Single Molecule Cluster Analysis (SiMCAn)^26,27^ of N=673 Hidden Markov Model (HMM) idealized smFRET traces from four experimental conditions (Fig. 2a-d): 5 mM Mg^2+^, 5 mM Mg^2+^ plus 400 nM RuvC, 5 mM Ca^2+^, and 5 mM Ca^2+^ plus 400 nM RuvC. Using a hierarchical cluster tree, we grouped traces with similar FRET states and kinetics into four clusters. We observed that 43% of the traces from the Mg^2+^/RuvC condition show distinct dynamic features, suggesting that they represent the ∼40% RuvC-bound, cleavage-enabled molecules; we therefore termed this cluster “R”. By contrast, the remaining clusters, termed S1, S2, and S3, showed protein-free, salt-condition-like *iso-I*↔*iso-II* dynamics and dominate in the absence of RuvC and/or presence of Ca^2+^ (Fig. 2f, g; Supplementary Fig. 5). Notably, a full 84% of all traces in the R-cluster arise from the Mg^2+^/RuvC condition, further supporting a unique behavior associated with the binding of both Mg^2+^ and RuvC. Accordingly, the FRET probability distribution of R-cluster traces populates the three conformations *iso-I* (high-FRET), Op (mid-FRET) and *iso-II* (low-FRET), with enrichment of Op from 23% to 50% unique to the Mg^2+^/RuvC condition (compare Figs. 2h and 2b, Supplementary Fig. 5c-d). The existence of *iso-I* and *iso-II* in these traces suggests that both conformers can be bound by RuvC in sequence and isoform-independent manner, and with unperturbed interconversion dynamics, consistent with prior observations of the PD state of RuvC^11^.

A transition occupancy density plot (TODP, Fig. 2i, Supplementary Fig. 5e) generated from over 900 transitions of RuvC bound R-cluster traces revealed three rapid, reversible conformational transitions: *iso-II*_*R*_↔*iso-I*_*R*_ (low↔high-FRET; 21%), *iso-II*_*R*_↔*Op*_*R*_ (low↔mid-FRET; 57%), and *iso-I*_*R*_↔*Op*_*R*_ (high↔mid-FRET; 22%). Among individual traces featuring the cleavage-enabled Op state, we observed three different classes: with only pairwise *iso-II*_*R*_↔*Op*_*R*_ transitions (45%), with only pairwise *iso-I*_*R*_↔*Op*_*R*_ transitions (10%), and with a slow interconversion from one to the other pairwise transition behaviors (35% “mixed” traces; Fig. 2e, Supplementary Fig.6). To further probe this interconversion, we segmented the mixed traces into their early and late phases and performed a second layer of SiMCAn. As expected, the early and late trace segments belong to distinct clusters (Supplementary Fig. 7a), for which TODPs revealed that the early and late phases preferably show *iso-I*_*R*_↔*Op*_*R*_ and *iso-II*_*R*_↔*Op*_*R*_ behavior, respectively (Fig. 2j). That is, we observe a strong propensity (>90% probability) of the mixed traces to start from *iso-I*_*R*_↔*Op*_*R*_ behavior and end with *iso-II*_*R*_↔*Op*_*R*_ behavior (Fig. 2k). Consistent with this finding, we observe an increase in traces showing the *iso-I*_*R*_↔*Op*_*R*_ to *iso-II*_*R*_↔*Op*_*R*_ interconversion from 20% to 32% when collecting traces at later time points (Supplementary Fig.7b). Significantly, Fano factor calculations reveal that, while the early *iso-I*_*R*_↔*Op*_*R*_ transitions are non-randomly (non-Poisson) distributed, the late-phase *iso-II*_*R*_↔*Op*_*R*_ transitions are random (Fig. 2l), providing direct evidence that the early phase still allows for PD-state mediated HJ branch migration, whereas the late phase does not.

To contextualize the complex HJ conformational isomerization we discovered upon RuvC binding under cleavage-enabled conditions, it is instructive to inspect the available crystal structures of the RuvC-HJ complex (Fig. 3). More than twelve amino acids from each monomer of RuvC form redundant contacts with the junction^6^, allowing the resolvase to remain loosely associated in the PD state so that the HJ can transition between isomers *iso-I* and *iso-II* as well as branch migrate (Fig. 3)^11^. Once the cognate cleavage sequence is approaching the enzyme’s catalytic core near the junction via this rapid sampling, charged amino acids such as Arg76 in the core of RuvC stabilize the G-C base of the consensus cleavage sequence via amino acid-nucleo base cation-pi interactions^6^. This positions the thymine residues of the cleavage sequence in hydrophobic pockets of the catalytic sites^3,8^ for HJ resolution. We propose that, once the cleavage-competent *iso-II* conformation is recognized, it snap-locks into place as evidenced by the near-irreversible switch from the early *iso-I*_*R*_↔*Op*_*R*_ to the late *iso-II*_*R*_↔*Op*_*R*_ pairwise transition behavior, positioning all RuvC-HJ contacts for catalysis (Fig. 3). At this stage, both transitions to *iso-I* and branch migration are inhibited, while the cleavage-competent *iso-II* conformation is snap-locked, ready for Mg^2+^-mediated catalysis. This model explains how the prototypic bacterial resolvase RuvC selects two of four possible HJ strands for cleavage. The RuvC dimer first recognizes and binds a HJ purely topologically, independent of sequence and conformational isomerization, then distorts the HJ to a partially open (Op) form^10^ while allowing it to remain dynamic by adopting its PD state. RuvC thus can utilize the intrinsic HJ dynamics through rapid conformational sampling and branch migration until it detects the cognate sequence in the continuous strand (here, of isomer *iso-II*), upon which amino acid-nucleo base interactions act as constraints to stabilize a cleavage-competent conformation through a near-irreversible conformational switch to *iso-II*_*R*_↔*Op*_*R*_, disallowing the cleavage-incompetent *iso-I* at minimal energetic cost. This constrained, snap-locked state is the basis for specificity during RuvC mediated cleavage, leading to a targeted set of desired recombination end products.

**Figure 3.**
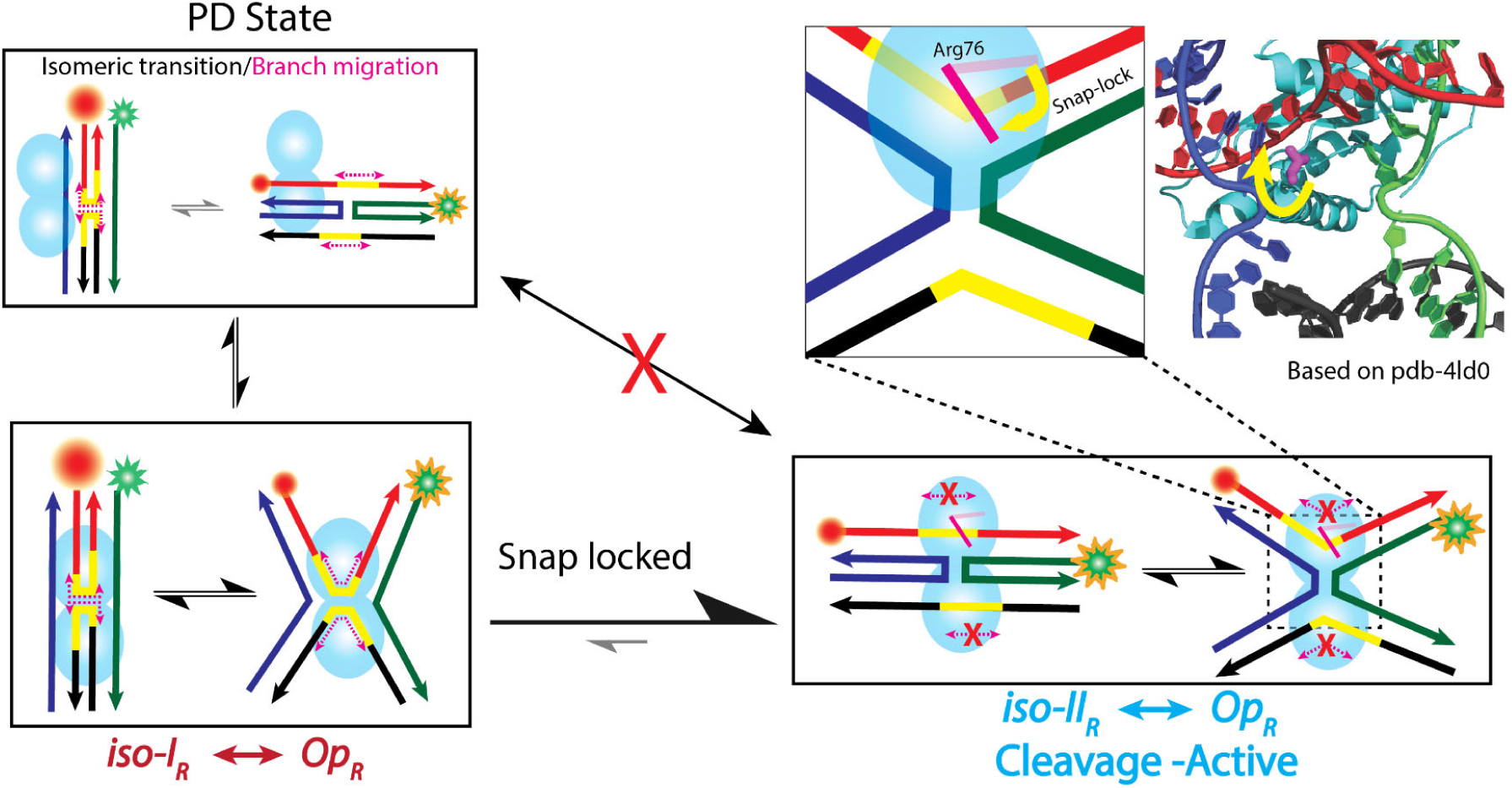
Proposed model for RuvC-mediated, site-specific Holliday junction resolution. When RuvC binds to *iso-I*, it forms an Open state (Op) where the cleavage sequence forms a shallow angle. RuvC allows the HJ junction to remain dynamic by partially dissociating from it (PD state). Isomeric transitions and branch migration are both allowed at this stage. Once a suitable cognate sequence is recognized, a (near-)irreversible transition leads to snap-locking of *iso-II* into the active site of the resolvase via amino acid-nucleo base interactions such as through Arg76, wherein the cleavage site can only still adopt the wider angle of a now cleavage-active Op state. Other conformational transitions such as to the PD state are suppressed.

In conclusion, we here have revealed the basis of RuvC’s unique combination of topological and sequence specificity, which may open exciting possibilities for antimicrobial therapy^28,29^ since sequence selectivity distinguishes RuvC from many eukaryotic/mammalian HJ resolvases^30–34^. Additionally, we anticipate that our quantitative single molecule cluster analysis will serve as a powerful tool to unravel the differentiating mechanisms of other DNA binding and processing enzymes.

## Methods

Methods, including statements of data availability and any associated accession codes and references, are available at …

## Acknowledgements

This work was supported by a NSF Award DMR-1607854 (to S.G.-T.) and NIH grant 2R01 GM062357 (to N.G.W.). We thank Dr. Alexander Johnson-Buck for careful reading of the manuscript and thoughtful suggestions.

## Author contributions

S.R., N.P., and N.G.W. designed the study and wrote the manuscript.

S.R. and N.P. performed all experiments and analyzed the data

S.R. and N.P. contributed equally.

## Additional information

**Supplementary information** is available for this paper at …

**Reprints and permissions information** is available at www.nature.com/reprints.

**Correspondence and requests for materials** should be addressed to N.G.W.

